# HF-125, a first-in-class computer-modeled novel inhibitor of Tribbles 2, for therapy of enzalutamide resistant, neuroendocrine prostate cancer

**DOI:** 10.64898/2026.04.27.720588

**Authors:** Sougata Ghosh Chowdhury, Pritam Biswas, Jitender Monga, Steve Brown, Craig Rogers, Jagadananda Ghosh

**Author notes:** ***Full name, mailing address, and email address of the Corresponding author:*** Jagadananda Ghosh, Department of Urology, Henry Ford Health System, 1 Ford Place, 2D, Detroit, MI 48202, USA.

## Abstract

Second generation antiandrogens, such as enzalutamide, are commonly prescribed to treat advanced prostate cancer. However, enzalutamide resistant prostate cancer (ERPC) invariably develops with more aggressive features. Management of ERPC is extremely difficult not only because available therapies cannot effectively eliminate ERPC cells but also due to enhanced growth and highly metastatic features in ERPC cells to invade distant organs. This problem is amplified by the lack of proper knowledge about suitable molecular targets in ERPC cells. Recently, we reported that Tribbles 2 (TRIB2), is overexpressed in ERPC cells and tumors and inhibition of TRIB2 kills ERPC cells via apoptosis. TRIB2 enhances cancer cell growth and invasion and confers resistance to enzalutamide, while inhibition of TRIB2 resensitizes the resistant cells. Interestingly, TRIB2 induces neuroendocrine (NE) features in ERPC cells and both the *de novo* and treatment-emergent NEPC cells consistently overexpress TRIB2. Though TRIB2 has emerged as a promising target, suitable inhibitors are not commercially available for clinical use. We used artificial intelligence (AI)-based molecular modeling to design a series of small chemical entities with potentially high specificity to bind with TRIB2. Extensive virtual screening and molecular editing yielded few highly selective and potent agents to inhibit TRIB2 and kill ERPC-NE/NEPC cells. One such compound (HF-125) directly binds to destabilize and degrade TRIB2 proteins involving proteasomes and is effective both *in vitro* and *in vivo*. Based on these findings, HF-125 emerges as a promising novel agent for development of a new therapeutic strategy for ERPC-NE/NEPC types of aggressive prostate cancer.

## Introduction

Projected diagnosis of ∼313,780 and death of ∼35,770 men in the year 2025 makes prostate cancer the most common form of malignancy and second leading cause of cancer-related deaths in American men **(1)**. Advanced prostate cancer is commonly treated with second generation androgen signaling blockers, such as enzalutamide, which directly binds and inhibits the androgen receptor function **(2-7)**. However, even after initial good response enzalutamide-resistant prostate cancer (ERPC) invariably develops which is lethal **(8-10)**. ERPC is not curable primarily because of the failure of available chemotherapeutic regimen to effectively kill the resistant cells. Moreover, lack of proper molecular understanding about mechanisms that are critical for survival of ERPC cells is delaying development of novel effective agents to selectively attack ERPC cells. Uncontrolled growth of aggressive ERPC cells eventually leads to widespread tumor metastasis causing excruciating pain and suffering to thousands of surviving patients. Thus, development of novel target-based agents to selectively eliminate ERPC cells is an area of utmost importance to delay progression of prostate cancer and prevent deaths. However, lack of proper knowledge about critical targets and pathways in the survival of ERPC cells is posing defeating challenges towards development of effective therapeutic agents. Hence, ERPC remains an enduring and debilitating health problem inflicting large numbers of patients and taking most of the lives lost due to prostate cancer.

To better understand the biology of ERPC, recently we developed an *in vitro* cell culture model and found gross overexpression of the pseudokinase, Tribbles 2 (TRIB2), in ERPC cells **(11, 12)**. In this model, parental enzalutamide sensitive prostate cancer cells were treated with gradually increasing doses of enzalutamide to mimic the standard clinical conditions in patients after prolonged therapy **(13,14)**. Overexpression of TRIB2 was also found in enzalutamide treatment resistant prostate tumor samples. Moreover, we found that forced overexpression of TRIB2 alone can convert sensitive cells to become resistant to enzalutamide treatment. Interestingly, inhibition of TRIB2 by shRNA re-sensitizes ERPC cells, suggesting that TRIB2 plays an important role in the resistance process and thus may emerge as a molecular target for development of a new targeted therapy for ERPC. An association of TRIB2 was also found with a range of other aggressive and drug-resistant types of cancers **(15-22)**. However, suitable small molecule agents are not commercially available for selective targeting of TRIB2 which is a requisite for proper clinical development. The main hurdle to develop agents to inhibit TRIB2 is its unique mechanism of action which involves protein-protein interactions followed by selective degradation of substrates. Because of the non-enzymatic nature of TRIB2 along with the absence of manageable deep pocket(s), development of quick and easy assay methods to analyze specific targeting agents to block the activity of TRIB2 protein-protein interaction is extremely difficult. Thus, in spite of being recognized as a *bona fide* target and promoter of therapeutic resistance in advanced prostate cancer, TRIB2 remains an elusive molecular target for developing an effective strategy to win over resistance to enzalutamide and other treatment modalities.

To overcome this problem, we used advanced computer programming to design a series of novel compounds bearing potential to strongly bind with the TRIB2 protein. After multiple rounds of ‘Design-Synthesis-Test’ cycles, we identified a new small molecule agent (called HF-125) which binds with TRIB2 and inhibits its activity with high selectivity and potency. Thermal shift assay revealed that HF-125 decreases the half maximal melting temperature (*Tm*) of recombinant pure TRIB2 protein, suggesting that HF-125 directly binds and destabilizes TRIB2 protein and primes it for enhanced degradation. Computerized prediction modeling suggested that HF-125 interacts with seven amino acid residues at the cleft region of TRIB2 protein, which is located closer to the N-terminus. In *in vitro* cellular assays we found that HF-125 effectively downregulates TRIB2 protein level and re-sensitizes enzalutamide-resistant prostate cancer cells. We also found that HF-125 downregulates neuroendocrine (NE) markers and induces apoptotic cell death. Moreover, HF-125 showed strong inhibition of ERPC tumor growth in SCID mice xenografts without any noticeable toxicity to general animal health. Thus, our in-silico drug design and development program, coupled with extensive characterization of compounds by structure-activity-relationship (SAR) studies, emerged as a powerful strategy to develop clinically competent novel compounds (such as HF-125) to effectively target TRIB2 both *in vitro* and *in vivo*.

## Results

### 1. High-throughput screening and computer-aided molecular editing identifies HF-125 as a novel agent to downregulate Tribbles 2

We found that HF-125 downregulates the protein level of TRIB2 in ERPC (LN-TRIB2) cell culture assays in a dose-dependent manner, while few of its analogs are ineffective **(Fig. 1A)**. Computer prediction modeling suggested that HF-125 binds to the cleft region of TRIB2 and presumably interacts with seven amino acid residues **(Fig. 1B-E)**. Thermal shift assay (TSA) showed that HF-125 decreased the half-maximal melting temperature (Tm) of purified recombinant TRIB2 protein, suggesting that HF-125 directly binds with TRIB2 protein and cellular thermal shift assay (CETSA) showed enhanced degradation of TRIB2 by HF-125 **(Fig. 1F-I)**. HF-125 dramatically degrades TRIB2 proteins in the presence of the cellular protein synthesis inhibitor, cycloheximide (CHX), and brings the half-life down to less than four hours **(Fig. 1J)**. Pretreatment of cells with the proteasome inhibitor (MG132) prevented HF-125-induced TRIB2 protein loss, suggesting involvement of proteasome activity in this process **(Fig. 1K)**.

**Figure 1.**
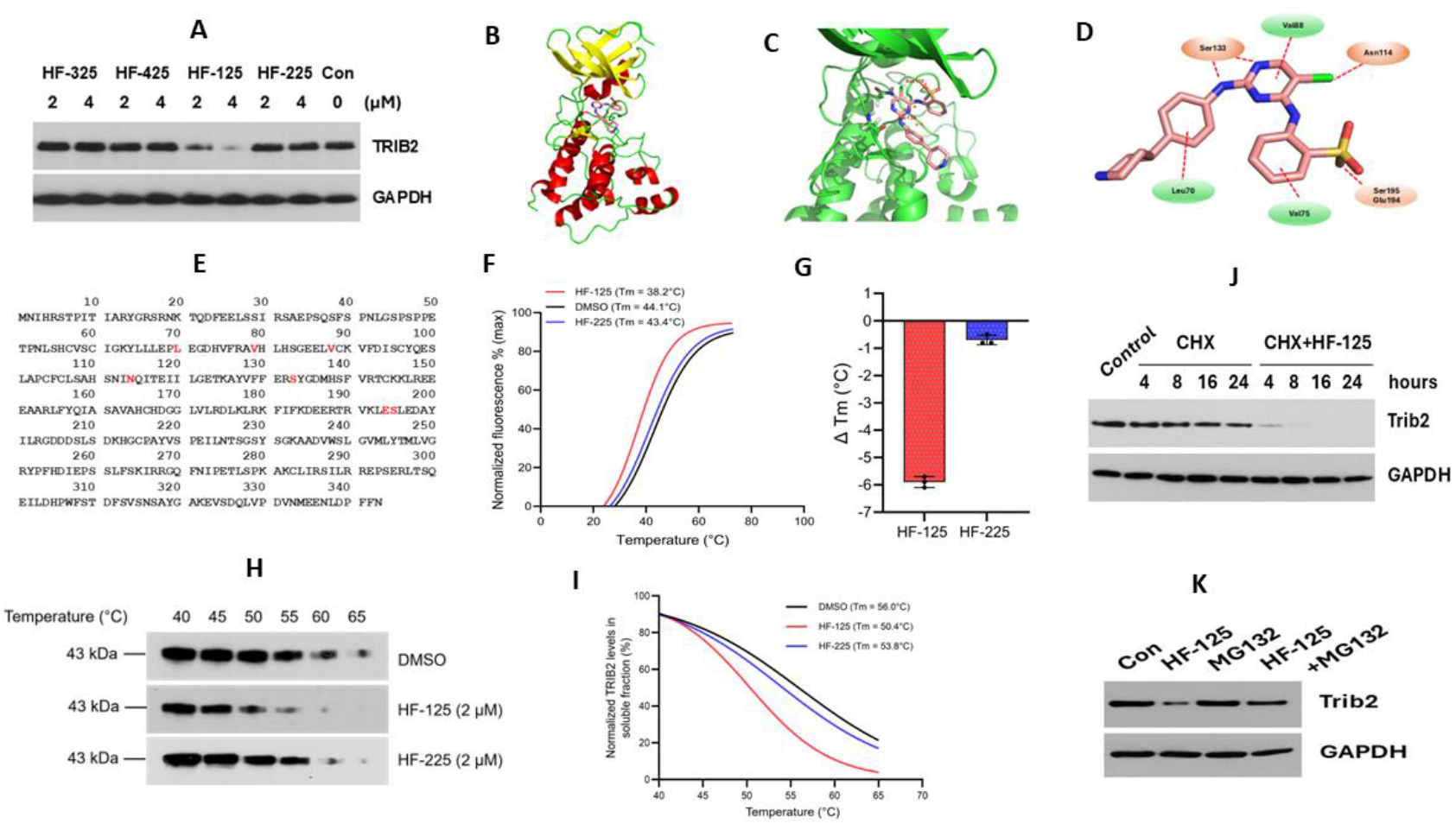
Characterization of HF-125. **A.** LN-TRIB2 cells were treated with four different compounds at the indicated concentrations. TRIB2 protein levels were determined by western blot analysis. GAPDH was used as an internal loading control. **B and C**. Molecular docking analysis of TRIB2 protein with HF-125 (ligand). **D**. Key binding pocket residues of TRIB2 participating in hydrogen bonding, hydrophobic interactions, and van der Waals interactions are highlighted. **E**. Amino acid sequence of TRIB2 showing potential binding site residues interacting with HF-125. **F**. Differential scanning fluorometry–based thermal shift analysis of purified protein TRIB2 revealed that HF-125 decreases the melting temperature (Tm), resulting in a leftward shift of the melting curve and indicating destabilization of TRIB2, whereas the other compounds HF-225 showed no significant change. ***Note:*** Chemical structures of HF-125 and HF-225 are provided in the Supplements section. **G**. Bar graph representing the change in melting temperature (ΔTm) of TRIB2 upon treatment with the indicated compounds, derived from differential scanning fluorometry. **H**. Cells treated with the indicated compounds (HF-125 & HF-225) or vehicles were subjected to thermal challenge at increasing temperatures. The soluble fraction of TRIB2 was analyzed by immunoblotting to evaluate compound-induced changes in protein stability. **I**. Band intensities from CETSA western blots were quantified and plotted as the relative soluble fraction of protein TRIB2 at each temperature for the indicated treatments. Data are presented as mean ± SD (n = 3 independent experiments). **J**. LN-TRIB2 cells were treated with cycloheximide (CHX) in the presence or absence of HF-125 and collected at four time points. TRIB2 protein levels were determined by immunoblotting to evaluate protein half-life, with GAPDH used as a loading control. **K**. Cells were treated either with HF-125 or MG132, or a combination of both. Controls were treated with vehicle (DMSO) only. TRIB2 protein expression level was assessed by western blot, with GAPDH used as a loading control.

### 2. HF-125 re-sensitizes enzalutamide-resistant prostate cancer cells

Since HF-125 effectively downregulates the protein level of TRIB2, we wanted to examine whether it can reverse enzalutamide resistance in prostate cancer cells which is contributed by high levels of TRIB2. We found that HF-125 effectively re-sensitizes both the treatment-induced enzalutamide resistant prostate cancer cells (LNCaP-ENR and MDA PCa-2b-ENR) as well as transfection/overexpression-induced enzalutamide resistant prostate cancer cells (LN-TRIB2 and MDA PCa-2b-TRIB2) by promoting DNA degradation and decreasing cell viability **(Fig. 2A-F)**.

**Figure 2.**
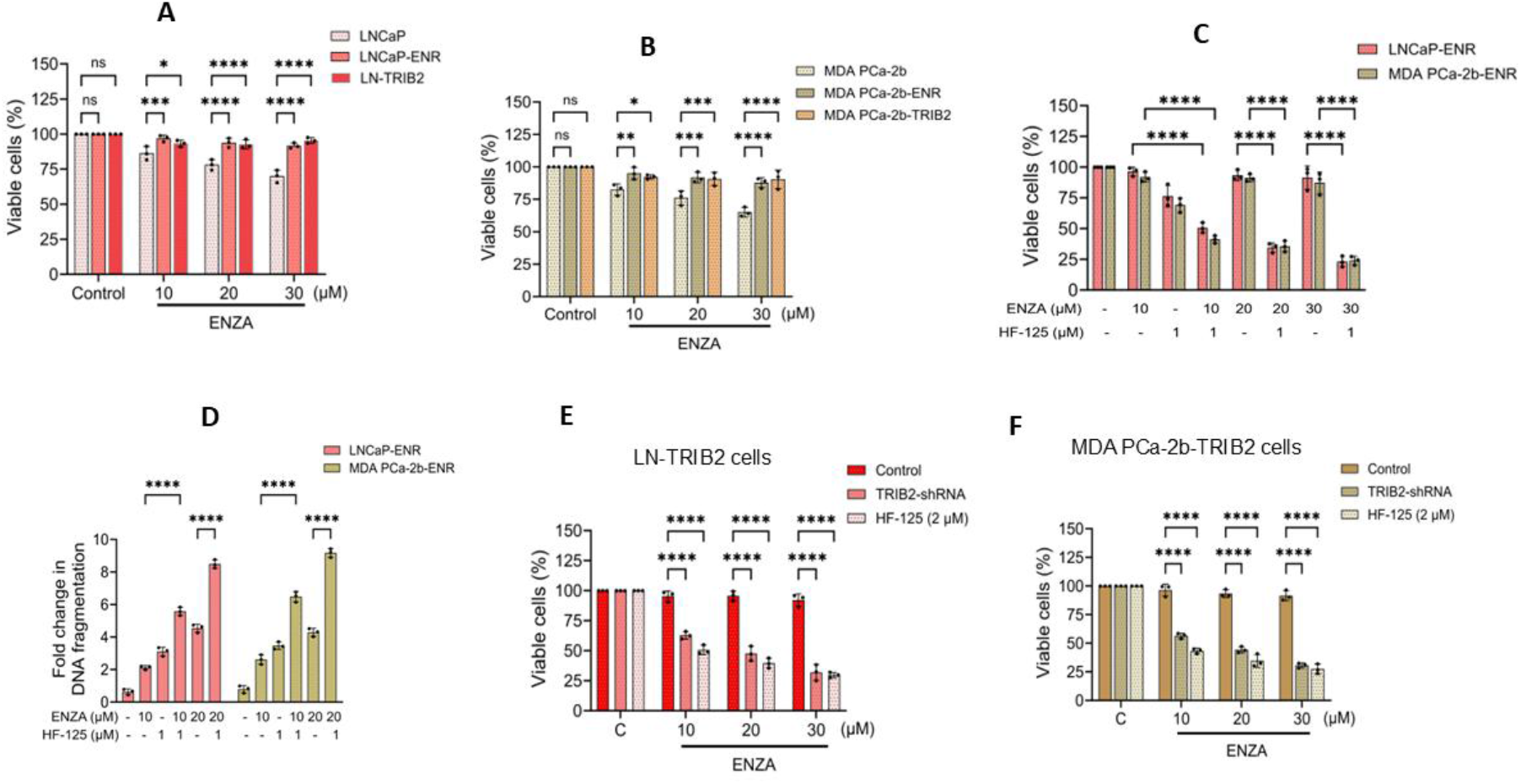
Re-sensitization of ERPC cells by HF-125. **A and B.** Dose-response analysis of enzalutamide in parental (LNCaP, MDA PCa-2b), enzalutamide-resistant (LNCaP-ENR, MDA PCa-2b-ENR) and TRIB2-overexpressed (LN-TRIB2, MDA PCa-2b-TRIB2) prostate cancer cell lines. **C**. Cell viability of enzalutamide-resistant (LNCaP-ENR, MDA PCa-2b-ENR) prostate cancer cells were evaluated following treatment with enzalutamide alone or in combination with the indicated drug HF-125. **D**. Enzalutamide-resistant cells were treated with enzalutamide in the presence or absence of HF-125, and fold change in DNA fragmentation was measured by ELISA to assess treatment-induced cell death. **E and F**. Enzalutamide-resistant (LNCaP-ENR, MDA-PCa-2b-ENR) prostate cancer cells were treated with HF-125 (2 μM) in the presence or absence of enzalutamide for 72 hours. In an independent experimental setup (n=3), Trib2 protein was silenced using shRNA instead of HF-125 for side-by-side comparison. Cell viability was measured using MTS/PES assays.

### 3. HF-125 downregulates neuroendocrine markers and kills androgen-resistant neuroendocrine prostate cancer cells

We observed that TRIB2 is grossly overexpressed in standard neuroendocrine prostate cancer (NEPC) cell lines and thus wanted to examine the effect of HF-125 on NEPC cell viability. We found that HF-125 decreases the viability of a series of NEPC cell lines in a dose-dependent manner **(Fig. 3A)**. Moreover, HF-125 dose-dependently decreased the protein levels of TRIB2 and several neuroendocrine marker proteins that are downstream targets of TRIB2 **(Fig. 3B-F)**. Interestingly, we found that androgen receptor (AR), which is downregulated by TRIB2, is upregulated by HF-125 treatment, presumably as a consequence of the inhibition/degradation of TRIB2.

**Figure 3.**
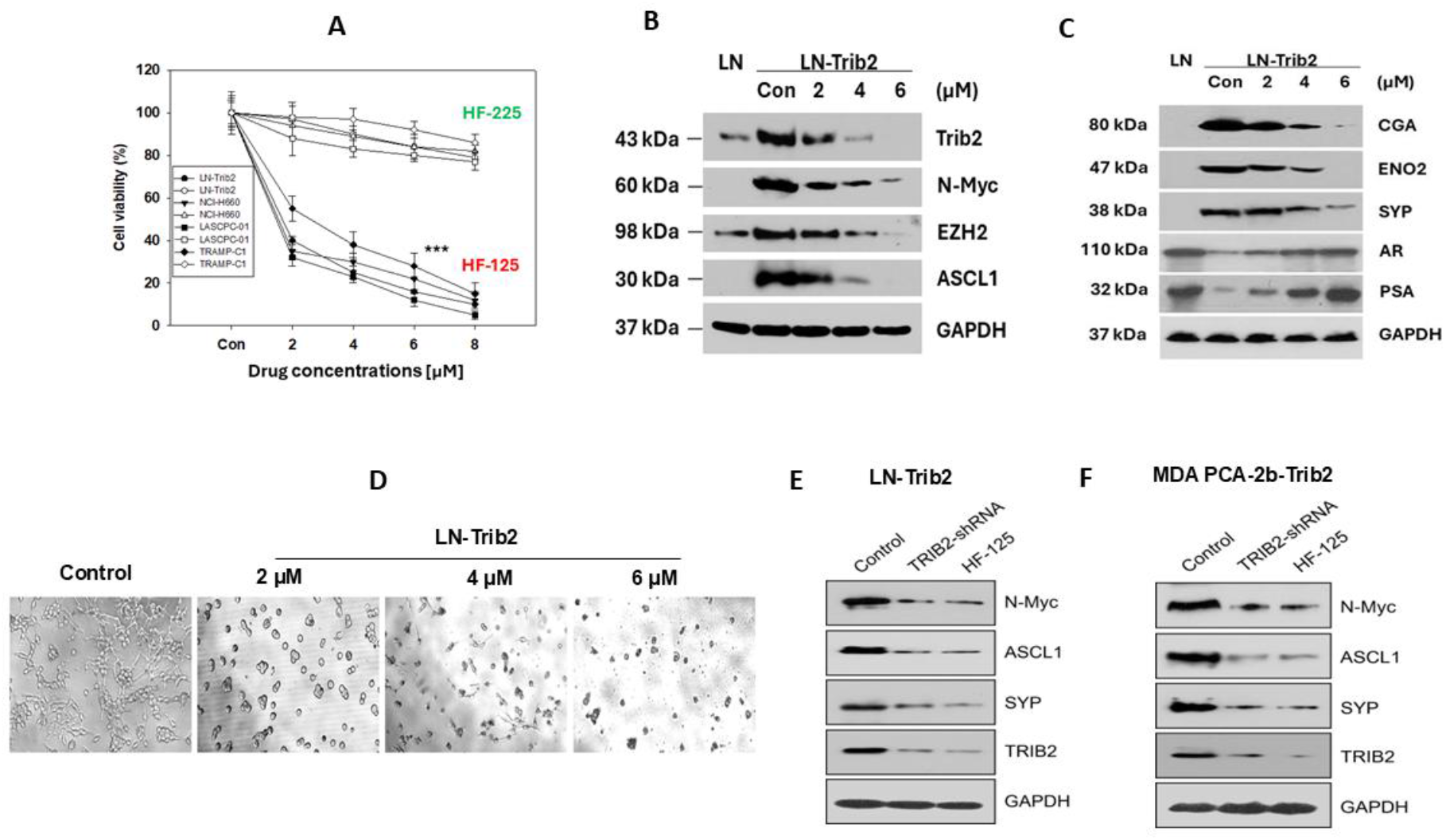
Molecular effects of HF-125 on ERPC-NE cells. **A.** About 3 × 10^3^ cells per well were seeded in 96-well plates in complete growth medium and exposed to varying concentrations of HF-125 or HF-225. Following 72 h incubation at 37°C in a humidified 5% CO_2_ incubator, cell viability was determined using the MTS/PES CellTiter assay (Promega). HF-125 significantly reduced cell viability, whereas HF-225 did not. **B and C**. LN-TRIB2 cells were seeded and treated with different concentrations of HF-125 for 48 h. Whole-cell lysates were collected, and proteins were resolved by SDS-PAGE. Expression levels of TRIB2 and other target proteins were assessed by western blot analysis. **D**. LN-TRIB2 cells were treated with varying doses of HF-125, and morphological changes were observed under phase contrast microscope. **E and F**. Western blot analysis demonstrates changes in TRIB2 and associated target proteins in TRIB2-overexpressed cells upon treatment with TRIB2-shRNA (1:10) or HF-125 at 2 μM.

### 4. HF-125 decreases colony growth and triggers programmed cell death (apoptosis) in ERPC-NE cells

Since we observed that HF-125 kills ERPC-NE/NEPC cells, we wanted to examine the mechanism involved in the cell death process. We found that HF-125 induces programmed cell death or apoptosis exerting cell cycle arrest presumably at the mitotic phase increasing G2M population **(Fig. 4A-D)**. We also found that HF-125 effectively decreases the colony formation and *in vitro* invasion of ERPC-NE cells in a dose-dependent manner **(Fig. 4E-H)**.

**Figure 4.**
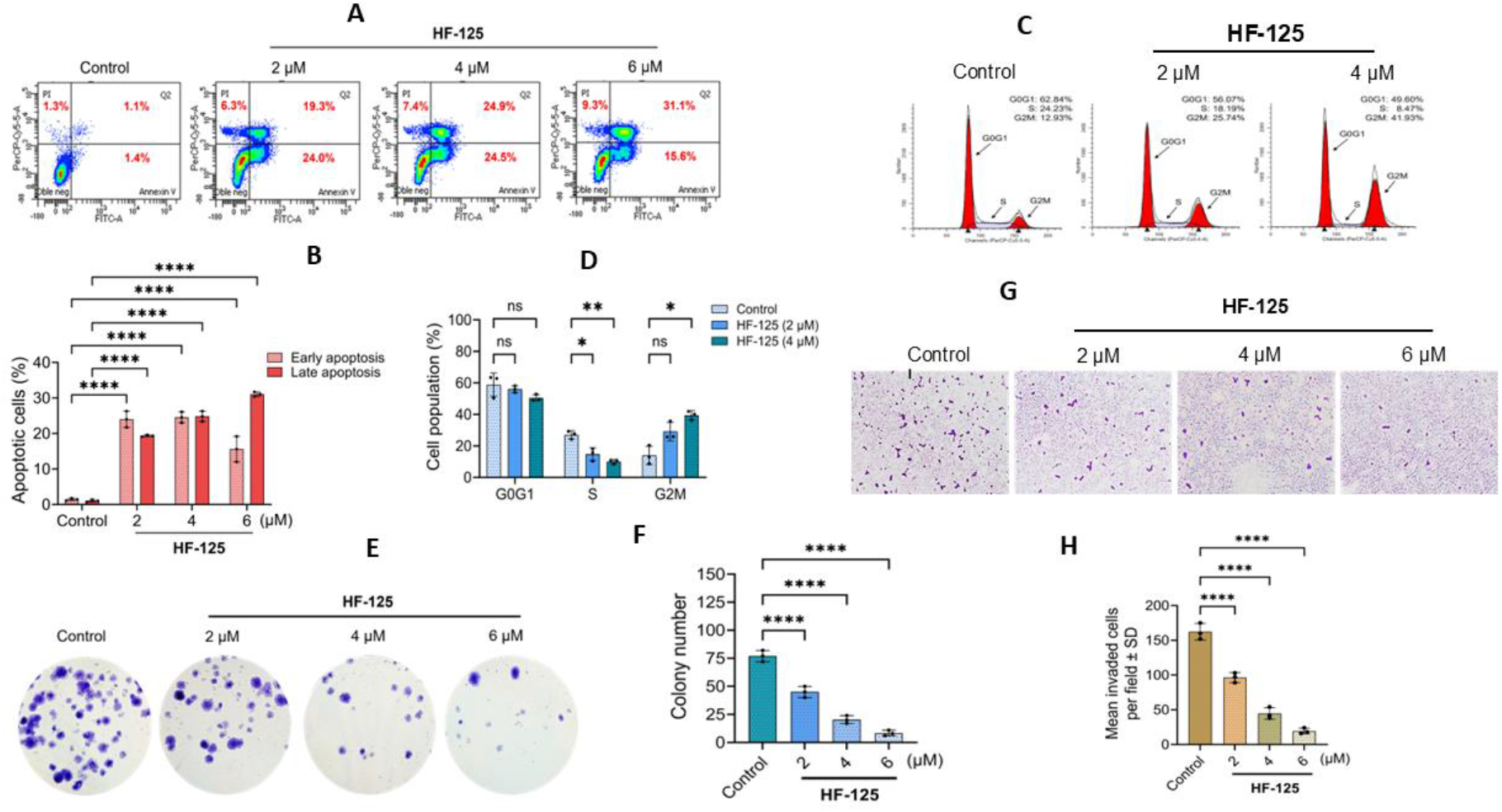
Effect of HF-125 on apoptosis, colony formation and invasion in ERPC-NE cells. **A.** LN-TRIB2 cells were treated with increasing concentrations of HF-125, and apoptosis was evaluated by detecting Annexin V binding using flow cytometry (FACS). **B**. Early and late apoptotic cell populations were quantified from three independent experiments and are presented as mean values. **C**. Vehicle (DMSO) or HF-125-treated cells were analyzed by flow cytometry to assess changes in cell cycle phase distribution. **D**. The percentage of cells in each phase of the cell cycle was quantified across different HF-125 doses to assess dose-dependent effects on cell cycle progression. **E**. LN-TRIB2 cells were treated with increasing concentrations of HF-125, and clonogenic potential was evaluated by colony formation assay. **F**. Quantitative analysis of colony numbers formed under increasing drug concentrations, demonstrating reduced long-term proliferative potential. **G**. *In vitro* invasion was measured by modified Boyden chamber assays demonstrating dose-dependent effect of HF-125 decreasing the invasive capacity of cells. **H**. Quantification of invaded cells is presented as mean values of each data point ± SE from three independent experiments, reflecting gradual loss in invasive potential in a dose-dependent manner.

### 5. HF-125 inhibits ERPC-NE tumor growth in SCID mice xenografts

Since HF-125 inhibits TRIB2 and kills ERPC-NE/NEPC cells, we wanted to examine whether HF-125 is effective *in vivo* to inhibit tumor growth. To investigate this, we subcutaneously implanted LN-TRIB2 cells into the flanks of SCID mice to develop tumors. Treatment started when tumors became measurable (around day 14 post tumor-implantation) and the mice were treated either with vehicle only (PBS) or HF-125 for four weeks **(Fig 5A)**. We observed that HF-125 at 40mg/kg/day strongly inhibits ERPC-NE tumor growth in xenografts and downregulates the protein levels of TRIB2 and targets without any overt toxicity to general animal health **(Fig. 5B-D)**. These findings suggest that HF-125 possesses excellent *vivo* efficacy and may turn out to be a suitable agent for therapy for ERPC-NE/NEPC type aggressive prostate cancer.

**Figure 5.**
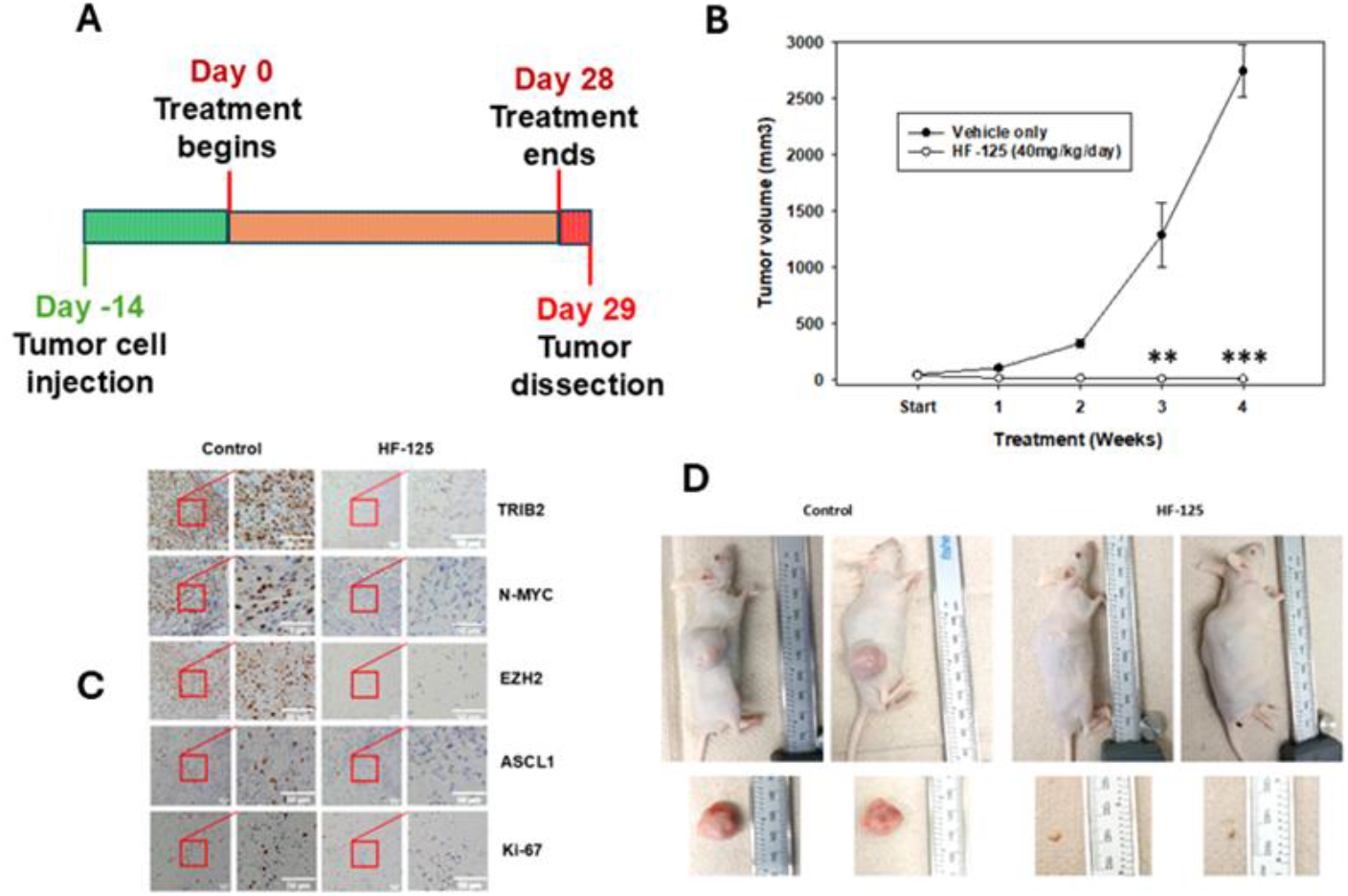
*In vivo* effect of HF-125 on ERPC-NE cell line xenografts. **A and B.** SCID-SHO mice were subcutaneously injected with LN-TRIB2 cells (4 × 10^6^ cells per mouse) to establish tumor xenografts and subsequently treated with HF-125 (40 mg/kg/day) or vehicle control via daily oral gavage for 4 weeks (n = 3). Tumor size and mice body weights were measured once per week. Tumor volumes were calculated using the formula ***TV*** = a × (b)^2^/2. *P* < 0.005. **C**. Tumor tissues from vehicle- and drug-treated mice were subjected to immunohistochemical (IHC) analysis to assess expression of selected protein markers. **D**. Representative control and HF-125-treated mice and tumors are shown.

### 6. HF-125 synergizes with enzalutamide to inhibit prostate cancer cells

Since HF-125 showed excellent TRIB2-inhibitory effects both *in vitro* and *in vivo*, we wanted to examine whether it can provide any benefit when applied together with enzalutamide. To evaluate the combinatorial effects of Enzalutamide and HF-125, cell viability was assessed following treatments with increasing concentrations of each drug alone or in combination in two prostate cancer cell lines (LNCaP and MDA PCa-2b). Interestingly, we observed that low doses of HF-125 and enzalutamide exert synergistic effects in decreasing the viability, invasion and induction of apoptosis **(Fig. 6A-F)** in LNCaP prostate cancer cells. Similar drug synergy was also observed in MDA PCa-2b cells **(Fig. 6G-I)**. Heatmaps of drug-induced inhibition of cell viability are shown in **Fig. 6A and 6G**. To further validate the synergistic effect, combination index (CI) values were calculated using the Chou–Talalay method **(28)**. CI versus fraction affected (Fa) plots revealed that the majority of data points fell below the line of additivity (CI = 1), confirming synergistic interactions between Enzalutamide and HF-125 in both cell lines **(Fig. 6B and 6H)**. *Note:* Strong synergistic effects (CI < 0.7) were observed with the two drugs. To quantitatively assess drug interactions, synergy was evaluated using the ZIP (Zero Interaction Potency) model. The ZIP synergy heatmaps demonstrated positive synergy scores across multiple dose combinations in both the cell lines **(Fig. 6C and 6I)**, indicating cooperative interaction between Enzalutamide and HF-125. Notably, the highest synergy scores were observed at intermediate concentrations, suggesting an optimal combinatorial window for maximal efficacy. The average ZIP synergy scores were positive in both cell lines, supporting a synergistic interaction rather than simple additivity. Changes in several apoptosis-related marker proteins were also observed both in LNCaP as well as MDA PCa-2b cells **(Fig. 6J, K)**. These findings suggest that HF-125 has excellent compatibility with enzalutamide and thus can be used together in combination to improve therapeutic index.

**Figure 6.**
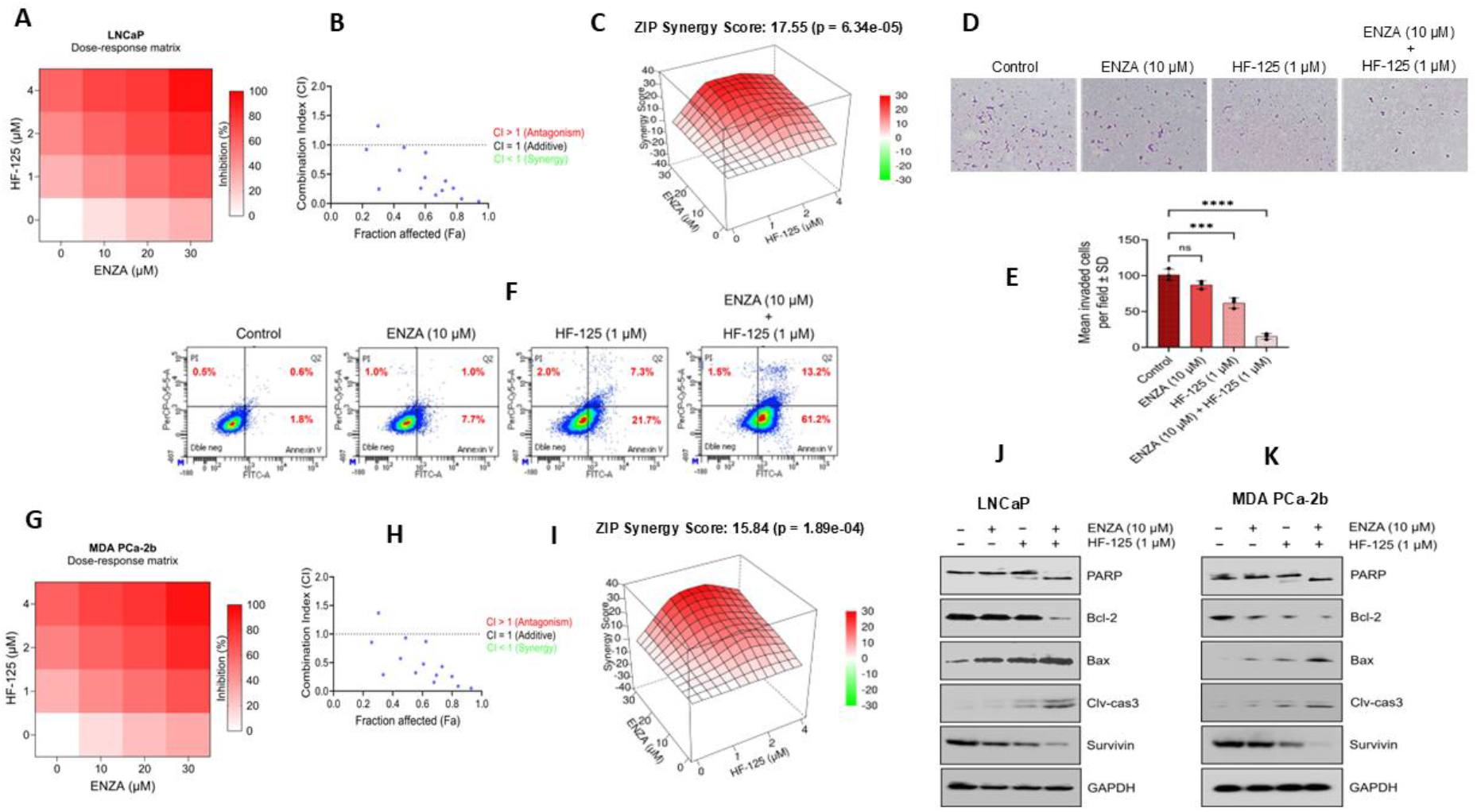
Synergistic effects of HF-125 and enzalutamide in prostate cancer cells. **A.** Heatmap showing cell viability (%) of LNCaP cells treated with increasing concentrations of Enzalutamide and HF-125 for 72 hours. Data normalized to vehicle-treated controls (100%). Color scale indicates relative viability. **B**. Combination index (CI) values were plotted using CompuSyn software for enzalutamide and HF-125 to evaluate synergistic interaction (CI < 1). **C**. The ZIP (Zero Interaction Potency) synergy scores were calculated using SynergyFinder, where positive scores indicate synergistic interaction **(38). D**. Representative invasion assay images showing control, individual drug treatments (Enzalutamide or HF-125), and combined drug treatment conditions. **E**. Quantification of invaded cells under control, individual drugs, and combined treatment conditions. **F**. Flow cytometric (FACS) analysis of apoptotic cell populations in control, single-drug (Enzalutamide or HF-125), and combined treatment groups. **G**. Heatmap showing cell viability (%) of MDA PCa-2b cells treated with increasing concentrations of Enzalutamide and HF-125 for 72 h. Data are normalized to vehicle-treated controls (100%). Color scale indicates relative viability. **H**. Combination index (CI) values were plotted against fraction affected (Fa) for enzalutamide and HF-125 to evaluate synergistic or antagonistic interactions. **I**. ZIP synergy scores were calculated to evaluate the interactions between enzalutamide and HF-125 in MDA PCa-2b cells. **J and K**. Western blot analysis of apoptotic proteins in LNCaP and MDA Pca-2b cell lines following treatment with drugs alone or in combination, indicating modulation of apoptosis-related pathways.

## Discussion

Enzalutamide is an FDA-approved popular drug which is frequently prescribed for advanced prostate cancer both in primary and metastatic settings. However, even after initial good response, enzalutamide resistant prostate cancer (ERPC) develops in almost every case and assumes a lethal phenotype. In fact, most of the deaths due to advanced prostate cancer happen because of the development of ERPC type of resistant cancer cells. Sadly, effective treatment options for patients inflicted with ERPC are extremely limited, primarily because of the lack of proper molecular understanding about critical targets to selectively attack and eliminate the resistant cancer cells without harming normal, non-cancer other body cells. Recently, we reported that the Tribbles 2 pseudokinase is grossly overexpressed in ERPC cells and plays a critical role in their survival. Thus, TRIB2 emerged as a promising new molecular target for ERPC therapy. Interestingly, we observed that overexpression of TRIB2 downregulates androgen receptor (AR) and transforms prostatic adenocarcinoma cells to develop neuroendocrine characteristics. Inhibition of TRIB2 reverses NE features and induces apoptotic cell death, suggesting a direct connection between TRIB2 and NE features. We also observed that overexpression of TRIB2 is a common feature in NEPC tumors and tumor-derived NEPC cell lines. Though TRIB2 appears to be an excellent molecular target for both treatment-emergent and de novo NEPC, suitable small molecule inhibitors are not commercially available for clinical use.

Development of targetable agents to inhibit TRIB2 is extremely difficult due to its non-enzymatic nature (a pseudokinase) and the absence of unique binding sites or manageable deep pockets in its three-dimensional structure. With the advent of advanced artificial intelligence (AI)-based molecular modeling, design and development of novel structure or ligand-based compounds became a reality. However, this approach bears high degrees of uncertainty and still needs wet lab confirmation. Using this approach, initially we screened a large library of compounds and obtained a number of initial hits potentially able to bind and inhibit TRIB2. After multiple rounds of derivatization and molecular editing following the principle of “Design-Synthesize-Test” cycles using both *in-silico* and *in vitro* cell culture testing, eventually we were able to generate few compounds with high selectivity and potency to inhibit TRIB2. One such compound was J1821-1629658, now called HF-125, which became the subject for further characterization because of its small size, excellent solubility profile and effect on TRIB2 proteins.

We found that HF-125 downregulates the protein level of TRIB2 in ERPC cells in a dose-dependent manner, while few of its congeners are ineffective. Computer prediction modeling suggests that HF-125 binds to the cleft region of TRIB2, presumably via non-covalent interactions with seven amino acid residues. Thermal shift and CETSA assays confirmed that HF-125 directly binds to destabilize and prime TRIB2 proteins for enhanced degradation involving proteasomes **(Fig. 1)**. Management of ERPC is extremely challenging because of the aggressive nature of ERPC cells, which are characterized by rapid growth and propensity to invade distant organs, including large bones. Loss of activity of enzalutamide in ERPC cells allow them to proliferate uncontrollably and metastasize which causes quicky deterioration of the health of prostate cancer patients. Interestingly, we found that HF-125 re-sensitizes the ERPC cells and effectively decreases their viability by inducing DNA degradation **(Fig. 2)**. This remarkable feature puts HF-125 into the clinical picture to be used even when standard FDA-approved androgen signaling blockade therapies become ineffective.

TRIB2, when overexpressed, not only makes prostate cancer cells resistant to defy enzalutamide treatment, but also it induces neuroendocrine features. Thus, our interest was to explore whether HF-125 by virtue of its inhibition of TRIB2, may inhibit the NEPC cells. By cellular and molecular analyses, we found that HF-125 effectively kills a range of NEPC cells and downregulates the protein levels of standard neuroendocrine markers (N-Myc, EZH2, ASCL1, CGA, ENO2, SYP) in a dose-dependent manner **(Fig. 3)**. One interesting finding is that HF-125 increased the protein levels of AR and its downstream target PSA. Presumably this happened due to the inhibition of TRIB2 by HF-125 and consequent removal of the repressive effects of TRIB2 on AR **(Fig. 3C)**. We were especially interested to know whether inhibition of TRIB2 by HF-125 can efficiently kill ERPC/NEPC cells because a majority of the drugs that are used in the clinic fail due to incomplete cell death, allowing residual cells to repopulate and continue tumor growth. We observed that HF-125 exerts cell cycle arrest, inhibits the colony-forming and invasive capability of ERPC-NE cells, and triggers apoptotic cell death as revealed by externalization of phosphatidylserine (reactive to Annexin V). These findings suggest that HF-125 may effectively debulk tumor load by *in vivo* by selective elimination of TRIB2-high aggressive cancer cells **(Fig. 4)**.

Excellent *in vitro* profile of HF-125 provoked us to examine the effects of HF-125 *in vivo*. Curiously, we observed that HF-125 showed dramatic inhibition of ERPC-NE tumor growth in SCID mice xenografts without any overt toxicity to the general health of treated animals. Moreover, HF-125 treatment showed strong downregulation in the protein levels of TRIB2, and related markers as evidenced by immunohistochemical analysis **(Fig. 5)**. An added benefit of new drug development is to see its compatibility and synergy with clinically relevant other drugs. Interestingly, we found that HF-125 exerts strong synergy when used in combination with enzalutamide at lower, sub-lethal doses, suggesting that these two drugs are highly compatible and thus, can be used together in combination **(Fig. 6)**. Altogether, our recent findings suggest that HF-125 may emerge as an effective novel therapeutic agent to control aggressive drug-resistant prostate cancer even with neuroendocrine characteristics.

Lineage plasticity and development of neuroendocrine features in therapy-resistant cells, including ERPC, is concerning and an alert to researchers to find novel avenues to circumvent the problem **(30-33)**. A role for lineage plasticity supported by cancer cell’s ability to reactivate the stemness program is getting increased visibility as a potentially new mechanism underlying development of resistance to therapy with various modalities. Prolonged inhibition of androgen receptor activity by second generation androgen signaling inhibitor drugs, such as enzalutamide is a prime example for induction of lineage plasticity and neuroendocrine differentiation **(34-37)**. However, the molecular underpinnings behind development and maintenance of neuroendocrine phenotypes are not fully understood and thus, development of suitable agents is delayed. Based on our recent work, it appears that TRIB2 plays a critical role in the biology of NEPC, and it can be targeted with appropriate agents to improve the current state of therapy. Based on chemical and biological properties, and the *in vitro* as well as *in vivo* activity, HF-125 emerges as a promising, target-based novel agent and ready for further characterization to be used as monotherapy or combination therapy for advanced, aggressive prostate cancer, including NEPC.

## Data Availability

All the data generated for this study are available within the article.

## Authors’ Contributions

- Conception, design, resources, funding acquisition: J. Ghosh
- Development of methodology: J. Ghosh, S. Ghosh Chowdhury, P. Biswas
- Analysis and interpretation of data: J. Ghosh, S. Ghosh Chowdhury, P. Biswas
- Writing, review, and/or revision of the manuscript: J. Ghosh, S. Ghosh Chowdhury
- Administrative, technical, or material support: J. Ghosh, S. Ghosh Chowdhury, P. Biswas, J. Monga, S. Brown, C. Rogers
- Supervision: J. Ghosh

## Acknowledgments

We acknowledge the help and cooperation from Dr. Patrick Eyers (University of Liverpool, UK) for sending us the 6x His-tagged TRIB2 plasmids and proteins. The authors deeply acknowledge the help and support from the Dykstra Foundation.

## Materials and Methods

### 1. Cell culture and reagents

Human prostate cancer cell lines (LNCaP, MDA PCa-2b) were obtained from the American Type Culture Collection (ATCC; Manassas, VA). The enzalutamide-resistant LNCaP-ENR and MDA PCa-2b-ENR cells were generated by treating with gradually increasing doses of enzalutamide over a 3 month-period. ERPC cells in culture were maintained in the presence of 30 µM enzalutamide. LNCaP-TRIB2 and MDA PCa-2b-TRIB2 cell lines were generated through stable overexpression of the full length human TRIB2 gene. Cells were cultured in RPMI-1640, or F12K supplemented with 10% fetal bovine serum (FBS) and 1% penicillin-streptomycin. Cell morphology was monitored regularly, and cultures were confirmed to be mycoplasma-free by PCR-based detection (Catalog #J66117; *A*lfa *A*esar, Tewksbury, MA).

### 2. Cell viability assay

Cells were seeded in 96-well tissue culture plates at a density of 3 × 10^3^ cells per well and incubated overnight to allow for attachment. Cells were treated with drugs and incubated for 72 hours in the CO2 incubator and cell viability was measured using the MTS/PES One Solution Cell Titer Assay (Promega Corp, Madison, WI) as described before **(23-25)**.

### 3. Microscopy

Cells were seeded at a density of 5 × 10^4^ per well in 6-well tissue culture plates in RPMI-1640 medium supplemented with 10% FBS. After 48 hours, the spent medium was replaced with 2 mL of fresh medium containing HF-125 or appropriate vehicle control. Phase-contrast images were captured at 20x magnification by a Keyence microscope using LJ-X8000 FileTransfer-MEA software.

### 4. Thermal shift assay (TSA)

Thermal shift assays were performed using an Applied Biosystems QuantStudio 6 Flex Real-Time PCR instrument following the Protein Thermal Shift Dye Kit protocol (Applied Biosystems). Pure TRIB2 protein was diluted in protein thermal shift buffer to a concentration of 5 μM and then incubated with HF-125 or HF-225 (2 µM) in a total reaction volume of 20 μl. The SYPRO Orange was used as a fluorescence probe.

### 5. Cellular thermal shift assay (CETSA)

Cells were plated and treated with HF-125 or HF-225 (2 μM) or vehicle for 2 h at 37 °C. Cells were harvested by scraping and resuspended in PBS supplemented with protease inhibitor cocktail. Cell suspensions were subjected to temperature gradient (40–65°C) for 3 min using a thermal cycler, followed by immediate cooling on ice for 3 min. Cells were lysed by three freeze–thaw cycles. Soluble proteins were resolved in SDS–PAGE and analyzed by Western blot. Band intensities were quantified using ImageJ and melting curves were generated by plotting normalized protein levels against temperature using GraphPad Prism software.

### 6. Western blot

Cells were seeded at a density of 3 × 10^5^ were plated in 60 mm plates, allowed to grow for 48 hours in the CO2 incubator and treated with drugs. Cells were harvested and lysed in lysis buffer (50 mM HEPES buffer, pH 7.4, 150 mM NaCl, 1 mM EDTA, 1 mM orthovanadate, 10mM sodium pyrophosphate, 10 mM sodium fluoride, 1% NP-40, and a cocktail of protease inhibitors). Proteins were separated by 10% or 12% SDS–PAGE and detected by Western blot using appropriate primary and secondary antibodies. Bands were visualized by ECL as described before **(26-28)**. Antibodies against TRIB2, SYP, CHGA, N-Myc, ASCL1, EZH2, AR, PSA, Bcl-2, Bax, PARP, Cleaved Caspase 3 and Ki-67 were from Cell Signaling Technology (CST) and GAPDH were from Santa Cruz Biotechnology (Santa Cruz, CA).

### 7. DNA-degradation assay

Cells (∼300,000) were plated in 60 mm diameter plates and allowed to grow for 48 h. The old medium was then replaced with fresh RPMI or F12K medium and then the cells were treated with drugs for 24 h. Drug-treated and control cells were lysed in lysis buffer for 60 min at 4°C and aliquots of lysates were used for measuring DNA-degradation to nucleosomal fragments using an ELISA kit from Roche (St. Louis, MO) as reported before **(26-28)**.

### 8. Invasion assay

*In vitro* invasion assay was done using Matrigel-coated Boyden transwell chambers from BD Biosciences. Transwells were soaked in 50 µl serum-free medium for 30 min at RT. Then ∼30,000 cells (in RPMI medium containing 0.1% BSA) were placed into the upper chambers and placed in 24-well tissue culture plates (one per well) on top of 500 µl RPMI medium containing 3% fetal bovine serum as chemo-attractant. Inhibitors were added directly to the medium and mixed. Plates were incubated at 37°C in the CO_2_ incubator for 16 h, processed as described before **(26-28)**, and observed under a Leica microscope at x200.

### 9. Computer modeling and drug design

Our basic approach was to find potential hits that selectively bind with Tribbles 2 protein preferably at the ATP competitive site. Using the AlphaFold2 and a proprietary computer-aided drug design program owned by MCSS (AU) we screened ∼3 million compounds, including about 8,000 compounds listed in the Drug Bank. Based on computerized homology modeling, we selected a list of twenty-five compounds carrying features with highest likelihood to bind strongly at the cleft region of the Trib2 molecule (closer to the N-terminus). After initial characterization by SAR studies, further modifications (derivatization) of selected hits were done by molecular editing and medicinal chemistry based on predicted values using multiple rounds of advanced more rigorous computer-based molecular dynamics program. Complete synthesis of selected compounds was done in batches by the medicinal chemistry group at JBL (IN) and supplied to us for *in vitro* cell culture and *in vivo* testing. Molecular mass and purity level (%) of each compound was provided by the manufacturer.

### 10. shRNA mediated gene knockdown

Knockdown of TRIB2 was done using lentiviral particles encoding shRNA obtained from the MISSION shRNA Lentiviral Transduction Particles (Sigma-Aldrich, St. Louis, MO, USA). Cells were plated and transduced with lentiviral particles at MOI 5 in the presence of 4 µg/mL Polybrene. Drug selection was done using puromycin (2 μg/mL).

### 11. Clonogenic assay

LN-TRIB2 cells were seeded in 6-well plates and treated with HF-125 or DMSO and cultured for 14 days. Cells were fixed in 1:7 Acetic acid/Methanol solution for 5 min at RT. After washing in PBS, crystal violet solution (0.5 %) was added and incubated for 20 min. Plates were washed with water and air-dried. Colonies were counted under microscope and calculated compared to vehicle control (100 % viable).

### 12. Flow cytometry

Apoptosis was measured using the Annexin V–FITC/Propidium Iodide (PI) apoptosis detection assay according to the manufacturer (BD Biosciences). Briefly, cells were seeded in 60 mm culture dishes and treated with the drug as indicated or vehicle control for 24 h. Following treatment, both floating and adherent cells were collected and washed twice with cold 1X PBS and resuspended in 1X binding buffer at a density of approximately 1 × 10^6^ cells/mL. Subsequently, 100 µL of the cell suspension was transferred into flow cytometry tubes and incubated with 5 µL Annexin V– FITC and 5 µL propidium iodide for 15 min at room temperature in the dark. After incubation, 400 µL of 1X binding buffer was added to each sample, and cells were analyzed by flow cytometry using a BD FACS Canto II. A minimum of 20,000 events were collected per sample. The percentages of early and late apoptotic cells were quantified for statistical analysis.

### 13. Cell cycle analysis

LN-TRIB2 cells were seeded in 60 mm diameter cell culture plates and treated with HF-125 for 24 h. Control was treated with the solvent (DMSO). Then, the cells were trypsinized and pelleted by centrifugation. Cell pellets were washed with 1X PBS and fixed with chilled 70% ethanol and kept overnight at −20°C. Cells were incubated for 10 min with 100 μg/mL of RNase A (Invitrogen) at 37°C, stained with 50 μg/mL propidium iodide, and cell cycle was analyzed by flow cytometry.

### 14. Drug combination assay

Cytotoxic effect of the individual drugs (enzalutamide or HF-125) or combinations were measured using MTS/PES assay. Drug synergy was determined using SynergyFinder 2.0 software. The synergistic interaction between drugs has a score greater than +10; an additive interaction has a score between -10 to +10; and an antagonistic interaction has a score of less than -10. Deviations between observed and expected responses with positive (red areas) and negative δ-values (green areas) indicate synergy and antagonism, respectively.

### 15. *In vivo* tumor xenografts

Animal studies were approved by the Institutional Animal Care and Use Committee (IACUC) and performed according to the institutional guidelines for animal care and handling. Exponentially growing LN-TRIB2 cells (4 × 10^6^ cells/mouse in 50% Matrigel in PBS) were subcutaneously injected into the right flanks of 6-week-old male SCID SHO mice (*n* = 4). When the tumors grew to ∼100 mm^**3**^, mice were randomly assigned and treated either with vehicle or HF-125 (40 mg/kg/day) via oral gavage for 4 weeks. Tumor size and mice body weights were measured once per week. Tumor growth was monitored by measuring sizes using digital slide-calipers and tumor volumes were calculated by the formula *TV* = a x (b)^2^/2.

### 16. Immunohistochemistry

Slides containing formalin-fixed paraffin-embedded sections were incubated at 60°C for 2 h, and antigen retrieval was carried out in EnVision FLEX target retrieval solution, Low pH (Agilent Dako, S236984–2) in a PT Link instrument (Agilent Dako, PT200). Slides were processed following methods as described before **(28)**. Images were captured at 20x magnification using a Leica microscope.

### 17. Statistical analysis

Statistical significance was assessed by two-way analysis of variance (ANOVA) or the two-tailed Students t-test and a value of <0.05 was defined as significant. Results are expressed as the mean value ± standard error of the mean (S.E.) and are described in figure legends when applied.

## Supplementary Data

**Supplementary Figure 1.**
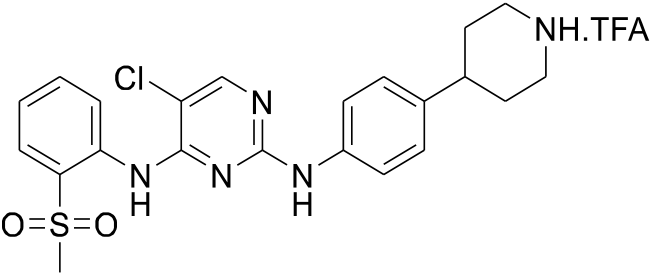
Chemical structure of HF-125

**Supplementary Figure 2.**
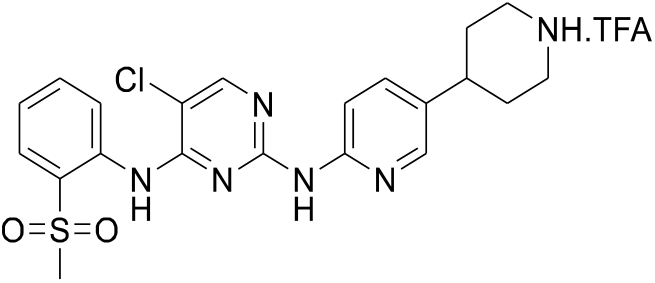
Chemical structure of HF-225

